# Demographic consequences of invasion by a native, controphic competitor to an insular bird population

**DOI:** 10.1101/182352

**Authors:** Kate M. Johnson, R.R. Germain, C.E. Tarwater, Peter Arcese

## Abstract

Species invasions and range shifts can lead to novel competitive interactions between historically resident and colonizing species, but the demographic consequences of such interactions remain controversial. We present results from field experiments and 45 yrs of demographic monitoring to test the hypothesis that the colonization of Mandarte Is., BC, Canada, by fox sparrows (*Passerella iliaca*) caused the long-term decline of the song sparrow (*Melospiza melodia*) population resident there. Several lines of evidence indicate that competition with fox sparrows for winter food reduced over-winter survival in juvenile song sparrows, enforcing population decline despite an increase in annual reproductive rate in song sparrows over the same period. In contrast, we found no evidence of interspecific competition for resources during the breeding season. Our results indicate that in the absence of a sufficient ecological or evolutionary shift in niche dimension, range expansions by dominant competitors have the potential to cause the extirpation of historically resident species when competitive interactions between them are strong and resources not equitably partitioned.

## INTRODUCTION

Exotic species are well-known to affect community composition via competitive, predatory and pathogenic interactions, especially on islands (Reaser et al. 2007; Dhondt 2012; Doherty et al. 2016). However, despite many native species undergoing range shifts linked to climate and land use change (e.g., Parmesan 2006; Early and Sax 2014; Krosby et al. 2015; Elmhagen et al. 2015), relatively little is known about the demographic impacts of these colonists on historically extant species (Davis and Shaw 2001; Loarie et al. 2008; Sorte et al. 2010; Rodewald and Arcese 2016). Theory suggests that the response of native species to controphic colonists will depend on their overlap in resource use, the demographic effects of resource limitation, the time-frame over which competitive exclusion might occur relative to the rate at which native species can adapt via ecological or evolutionary shifts in niche dimension, and the spatial scale examined (Shea and Chesson 2002; Davis 2003; Gurevitch and Padilla 2004; Reaser et al. 2007; Sax et al. 2007; Bennett et al. 2012; Dhondt 2012; Stuart et al. 2014). Although examples of competitive exclusion by colonist species remain rare, they are thought to be most likely to occur where environmental heterogeneity is low, such as small islands or isolated water bodies (e.g., Chesson 2000; Davies et al. 2005; Melbourne et al. 2007; MacDougall et al. 2009). Given the potential for novel competitive interactions to shape species demography and community composition (Simberloff 2005; Reaser et al. 2007), we used a 45 yr study of an island song sparrow (*Melospiza melodia*) population to ask how the colonisation and expansion of a colonist, controphic competitor, the fox sparrow (*Paserella illiaca*) has affected its demography.

Although described as migratory throughout its range in North America (Weckstein et al. 2002), fox sparrows established resident populations in the San Juan and Gulf Islands of western North America in the latter half of the 20^th^ century, where they now survive and reproduce at rates consistent with rapid population growth (Visty et al. 2017). Because song and fox sparrows are territorial, very similar in life history, but differ in size, we used field experiments and demographic analyses to test for evidence of interspecific competition between song sparrows and this colonist following Dhondt (2012) and Jankowski et al. (2010).

Specifically, we first used a life-table response experiment and 45 years of demographic data to identify vital rates contributing most to song sparrow population growth over time, and to test if those rates varied with fox sparrow abundance. We next tested for interspecific competition for breeding habitat by conducting simulated territorial intrusions to quantify interference competition. Because exploitative competition might reduce breeding habitat quality even in the absence of interspecific territoriality, we also tested for long-term declines in site quality following Germain and Arcese (2014). Last, we tested for evidence of interspecific competition for access to winter food by measuring diet overlap and inter-specific dominance.

## METHODS

### Study system

Mandarte Is. is a *c.* 6 ha islet in southwestern BC, Canada, where a resident, individually-banded song sparrow population was monitored from 1960–63 and 1975–2016 (Tompa 1963; Germain et al. 2015). Song sparrows are a *c.* 24 g passerine that occur over much of North America at densities of ∼1–9 pairs/ha (Arcese et al. 2002). On Mandarte Is. song sparrows lay 2–5 eggs in 1-7 open-cup nests annually (Arcese et al. 1992) and defend 200–5000 m^2^ territories year-round (Arcese 1989). From April – July 1960–63, 1975-79, and 1981-2016, the territorial status and breeding activity of all song sparrows was monitored at least weekly to locate all nests annually. All nestlings were colour-banded, followed to independence from parental care (∼24 days post-hatch) and their recruitment or disappearance from the population was recorded, which provided precise estimates of annual population size, age structure, and reproductive and survival rates (annual re-sighting probability >99%; Wilson et al. 2007). About one immigrant song sparrow settles on Mandarte Is. annually, but because that rate is low and about constant it is not considered here.

Fox sparrows are ∼19% larger than song sparrows in mass and linear traits and, like song sparrows are territorial, multi-brooded, open-cup nesters that feed mainly on seeds (winter) and insects (breeding; Weckstein et al. 2002, Visty et al. 2017). Fox sparrows are native to BC, but were absent from Mandarte Is. prior to 1975 (Tompa 1963; Drent et al. 1964), when they first bred there (J.N.M. Smith, pers. com.). Fox sparrows were counted systematically in 13 yrs from 1960-2016 by spot-mapping singing males, their mates and nest locations, and by mapping their territories in detail in 2010 and 2013–16 when up to 70% of adults were individually identified.

Rate of annual change in the number of female sparrows on the island in late April was estimated using a generalized linear model with year as a fixed effect (fox sparrow: Poisson distribution, log link; song sparrow, Gaussian distribution). We tested for temporal autocorrelation in the time series (finding none) and employed R 3.1.3 (R Core Team 2015) for all statistical analyses.

### Demographic rates

We identified the demographic vital rate contributing most to song sparrow population growth using a stage-structured life table response experiment (LTRE) to estimate the contribution of each vital rate to growth from 1975–2014. We used juvenile and adult age-classes for both sexes because adults differ modestly in survival and reproductive rate after reaching adulthood. Local juvenile survival was estimated as the proportion of offspring surviving from independence (day 24 post-hatch) to April 30 the next year. Adult survival was the proportion of individuals alive on April 30 in year *t* that survived to year *t* + 1. Reproductive rate was the mean number of independent young produced per female annually, excluding birds used in experiments (1979, n = 70; 1985, n = 87; 1988, n = 114; Arcese and Smith 1988; Smith et al. 2006). Juvenile survival was unknown in 1979 and 1980, reproductive rate unknown in 1980, and adult survival unknown in 1975.

The LTRE included a treatment matrix parameterized with juvenile and adult survival and adult reproductive rate in each of 37 yrs. Vital rates were arranged in 2 × 2 treatment matrices wherein the 1^st^ and 2^nd^ columns included juvenile and adult vital rates, respectively, and the first and second rows specified reproduction and survival rates, respectively. Because juveniles do not breed, the 1^st^ row of column one always equalled zero. Treatment matrices were compared to a 2 × 2 reference matrix of mean vital rate over all years to determine the contribution (*c*_*ij*_) of each vital rate to annual population growth (*cf* Caswell 1996) such that:

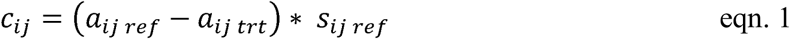

where *a*_*ij*_ is the (*i,j*) element of *a*, the reference (*ref*) or treatment (*trt*) matrix, and *s*_*ij*_ is the sensitivity of the reference matrix, indicating the impact of an absolute change in a vital rate on population growth (de Kroon et al. 1986; Caswell 1996). Analyses were implemented in the *popbio* package (Stubben and Milligan 2007) and trends in vital rate contribution estimated by linear model (Gaussian). Because the LTRE indicated that juvenile survival was a key vital rate, we tested for an effect (α ≤ 0.05) of fox sparrow abundance on juvenile song sparrow survival using a linear model (Gaussian distribution), with fox and song sparrow numbers as predictors.Song sparrow numbers was included because it was shown previously to predict juvenile survival in that species (Arcese et al. 1992). Estimates of fox sparrow population size and song sparrow juvenile survival were available for 11 years from 1960 to 2016.

### Competition for winter food

Dhondt (2012) identified space, nesting habitat and food as common limiting factors in bird communities. We tested for interspecific competition for winter food between fox and song sparrows using a seed preference experiment to estimate diet overlap, and two arena experiments to assess behavioral dominance in contests over food. We assessed the breadth of winter food available by characterising the type and abundance of seeds in soil, given that both species feed mainly on seeds in winter (Tompa 1963; Willson 1971; Arcese et al. 1992; Weckstein et al. 2002). To do so we excavated 250 ml of soil (10 × 15cm, 2cm deep) at 15 sites across the island in December 2013, extracting all seeds by sieve to estimate abundance by identity. Seeds recovered in soil included 64% blackberry (*Rubus armeniacus*), 17% Oregon grape (*Mahonia aquifolium*), 8% Nootka rose (*Rosa nootkensis*), 8% red elderberry (*Sambucus racemosa),* and 3% other (bitter cherry, *Prunus emarginata;* choke cherry, *P. virginiana;* snowberry, *Symphoricarpos albus; grape, Vitis sp.*) by volume.

We estimated fox and song sparrow preference for seeds in March 2015 by cleaning, then freezing blackberry, Nootka rose, elderberry, and snowberry seeds collected from fruits in summer 2014. We chose the seed types that were most abundant in soil samples with the exception of Oregon grape and cherry, which are ∼1.5× larger than all other seed types and are likely inedible to song and fox sparrows. Seeds were arranged by species in one of four 98cm^3^ circular depressions (‘cups’) in 60 × 12 × 3 cm plywood feeders, and rotated among cups in each trial to avoid location effects. Feeders were placed on the ground at 6 locations on the island used regularly by foraging fox and song sparrows. In each trial we recorded by video the fraction of time a visiting fox or song sparrow fed on each seed type (N=14 trials, including 50 visits by 6 different song sparrows and 9 different fox sparrows). A ‘visit’ comprised the time elapsed from when a focal bird picked up its first seed, to the time the focal bird’s lower mandible stopped moving after last seed was eaten, prior to leaving. Seed preference was estimated by recording the total time from when a focal bird picked up a first seed in cup *x*, to the point its lower mandible stopped moving before selecting a seed from a different cup or leaving. The proportion of total time spent feeding on each seed type during a visit was then used as the dependent variable in a generalized linear mixed model (GLMM, quasibinomial distribution, logit link). Each visit was numbered and included in the model as a random effect, as was fox or song sparrow identity. We used the glht function in the *multcomp* package (Hothorn et al. 2008) to assess statistical significance of all pairwise comparisons of fox or song sparrow with each seed species using Tukey contrasts for unequal groups (Tukey 1949; Kramer 1956).

We conducted arena experiments in October 2013 to assess interspecific dominance at winter food sources by piling 250ml of commercial bird seed at 5 sheltered locations across the island and using video cameras to record interactions. We recorded 19 fox and 19 song sparrows in 68 aggressive interactions. The wining individual stayed in the arena (datum = 1); the loser was chased (datum = 0; Arcese and Smith 1985). We tested the null expectation of equality among species using two GLMMs (binomial distributions, logit links), each including fox and song sparrow identity as random effects. In the first model, we estimated the displacement rate using maximum likelihood; in the second model, we tested the hypothesis of displacement rate equal to 0.5 for both species. Finding a significant effect of ‘species’ on displacement is consistent with a hypothesis of interspecific dominance. We replicated the arena experiment in March 2015 using 20 feeders (15 × 20cm plastic tray on ∼20cm stake; 250ml of commercial birdseed), distributed in sheltered sites across the island. We monitored, scored and analysed 31 interactions between 20 song and 16 fox sparrows.

### Competition for space

To test whether fox and song sparrows compete for breeding space, we first calculated the spatial overlap of song and fox sparrow territories in 2010, 2013, 2014 using ArcMap (ESRI 2011). We also conducted playback experiments prior to breeding in April 2014 to quantify the response of song sparrows to fox sparrow, song sparrow, and Swainson’s thrush (*Catharus ustulatus*) (a control species) following Jankowski et al. (2010). Playback experiments involved placing a taxidermic mount on an artificial perch at the center of 27 song sparrow territories, and playing species-appropriate song from a speaker placed below the mount. In each 12 min trial we recorded the closest approach of the territorial male and female song sparrow to the mounts, during: 2 min of pre-trial observation, 5 min of playback, and 5 min follow-up observation. All mounts were presented in random order and prepared in a neutral, perched position. Swainson’s thrushes are similar to fox sparrows in size but do not breed on Mandarte Is. Trials were conducted throughout the day but separated by ≥ 1 hour on focal and neighboring territories. Closest approach to the mount was used as a dependent variable to indicate aggression (*cf* Jankowski et al. 2010) and compared among mounts using a GLMM (negative binomial distribution, log link), and including the identity of focal male song sparrows as a random effect. Time of day (categorical effect: morning, <10am, n = 22; midday, 10–1pm, n = 27; afternoon, 1– 4pm, n = 6; evening, >4pm, n = 22), whether or not the focal male was singing prior to playback (1/0 fixed effect), and whether one or more neighbor males sang in response to playbacks were also recorded and included as covariates (1/0 fixed effect).

### Competition for nesting habitat

We followed Germain and Arcese (2014) to estimate site quality as the mean number of independent song sparrow young produced annually in each of 146 20 × 20 m grid cells distributed continuously over the island. Doing so allowed us to test the prediction that site quality for song sparrows declined as fox sparrow abundance increased over time, as expected given competition by song and fox sparrows for breeding resources. Specifically, we regressed site quality on year using a linear mixed model (year of study included as a fixed effect, grid cell identity as a random effect). We also tested for a decline in the mean number of independent young produced by female song sparrows over the study using a GLMM (negative binomial distribution, log link) with year as a fixed effect and female identity as a random effect to account for repeat observations across years.

## RESULTS

### Population size and demography

Female song sparrow population size varied widely over 45 years (range = 4–71, mean = 35.0 ± 17.1 SD), but declined on average (β = −0.60 ± 0.14 SE, *t*(43) = −4.19, p < 0.001, R^2^ = 0.29; figure 1). In contrast, fox sparrow abundance increased after 1975 (range = 1–30 females, β = 0.06 ± 0.01 SE, *z*(11) = 9.13, p < 0.001; figure 1).

**FIGURE 1.**
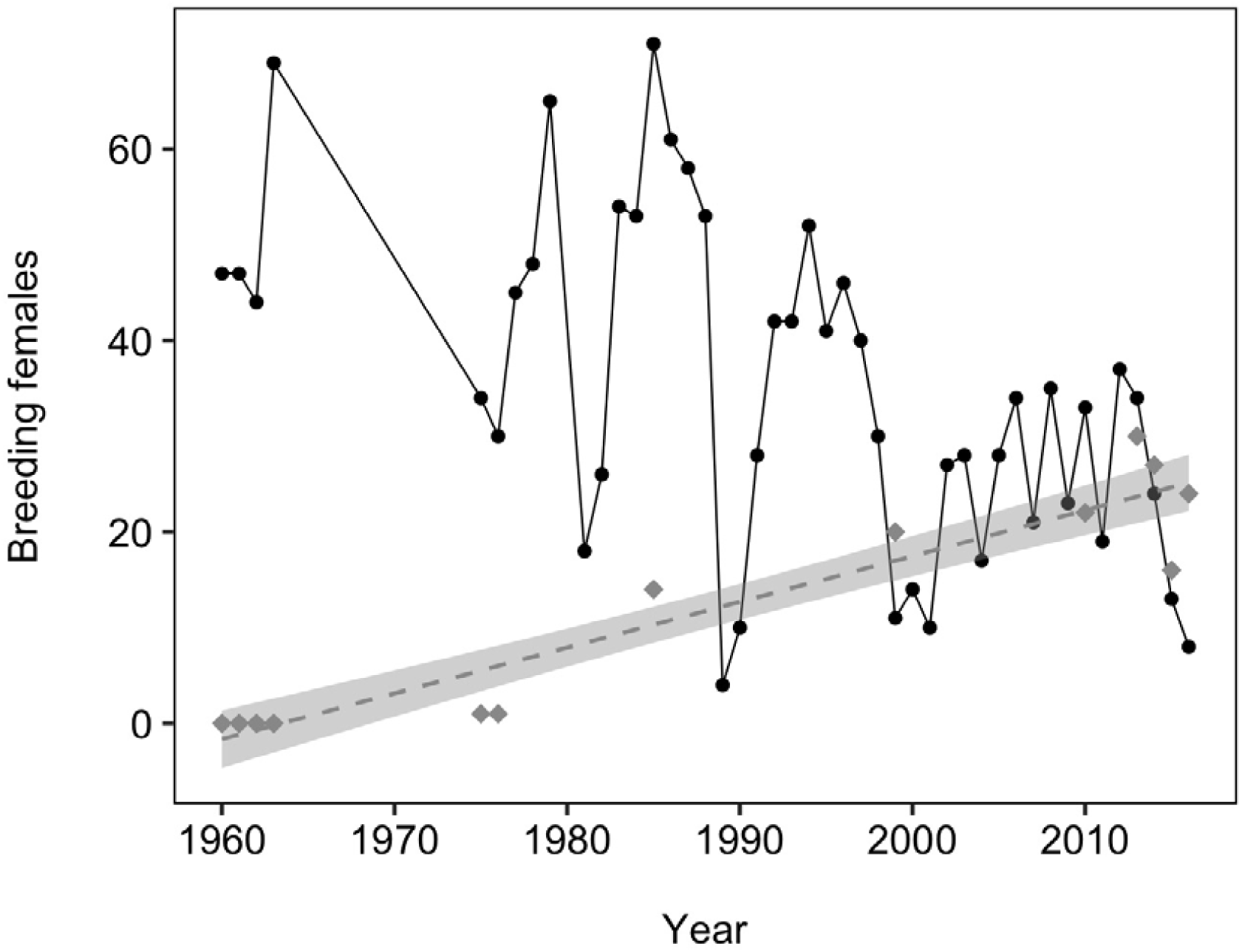
Number of breeding female song sparrows in 45 years from 1960–63 and 1975–2016 (black circles) and fox sparrow breeding females in 13 years from 1960–2016 (grey diamonds). Song sparrow population size has declined significantly over the study period while fox sparrow population size has increased. Shaded areas represent predicted values ± 1 SE.

Juvenile song sparrow survival also varied widely over the 37 yrs it was recorded from 1960 to 2016 (range = 0.04–0.88, mean = 0.37 ± 0.18 SD, *n*_yrs_ = 37; figure 2), as did adult annual survival (range = 0.07–0.88, mean = 0.59 ± 0.17 SD, *n*_yrs_ = 38), and annual reproductive rate (range = 1.10–6.90, mean = 3.25 ± 1.32 SD, *n*_yrs_ = 38). However, despite wide variation in song sparrow vital rates over time, when contribution (as determined from the LTRE) was regressed on year, juvenile survival was the only vital rate to increase in influence over time (β = −0.01 ± 0.004 SE, *t*(35) = −3.37, R^2^ = 0.25, p = 0.002; figure 3a). Variation in annual reproductive rate had no detectable effect on long-term change in song sparrow population size (*t*(35) = −1.29, p =0.21, R^2^ = 0.05; figure 3b). Similarly, the contributions of annual adult survival to change in population size were smaller and unrelated to long-term decline in song sparrow abundance (*t*(35) = 1.29, p = 0.20, R^2^ = 0.05; figure 3c).

**FIGURE 2:**
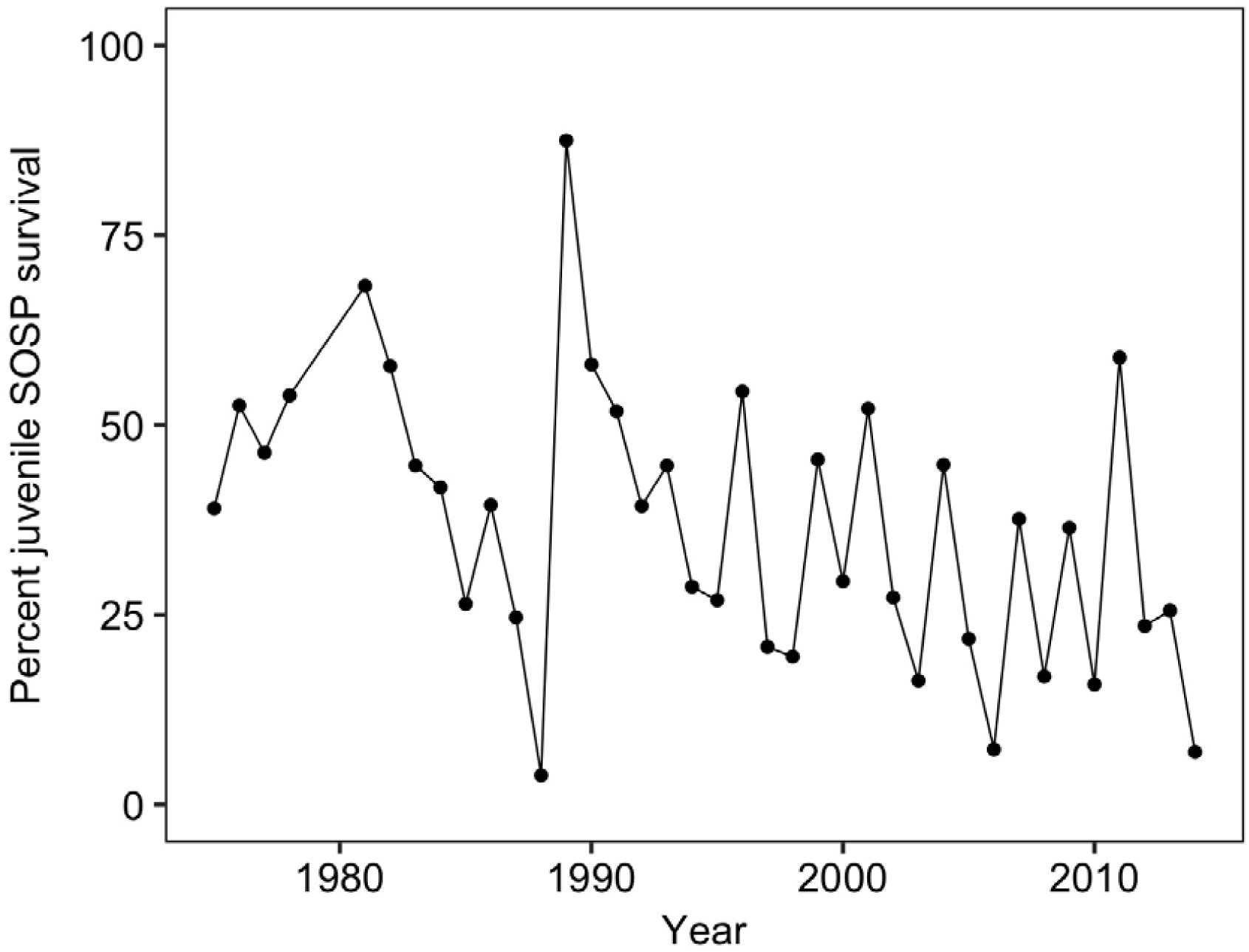
Percent of yearling male and female song sparrows surviving from the end of parental care (day 24 after hatching) to April 30th of the following year (juvenile survival) from 1975-2016 (excluding 1979 and 1980 when juvenile survival was unknown).

**FIGURE 3.**
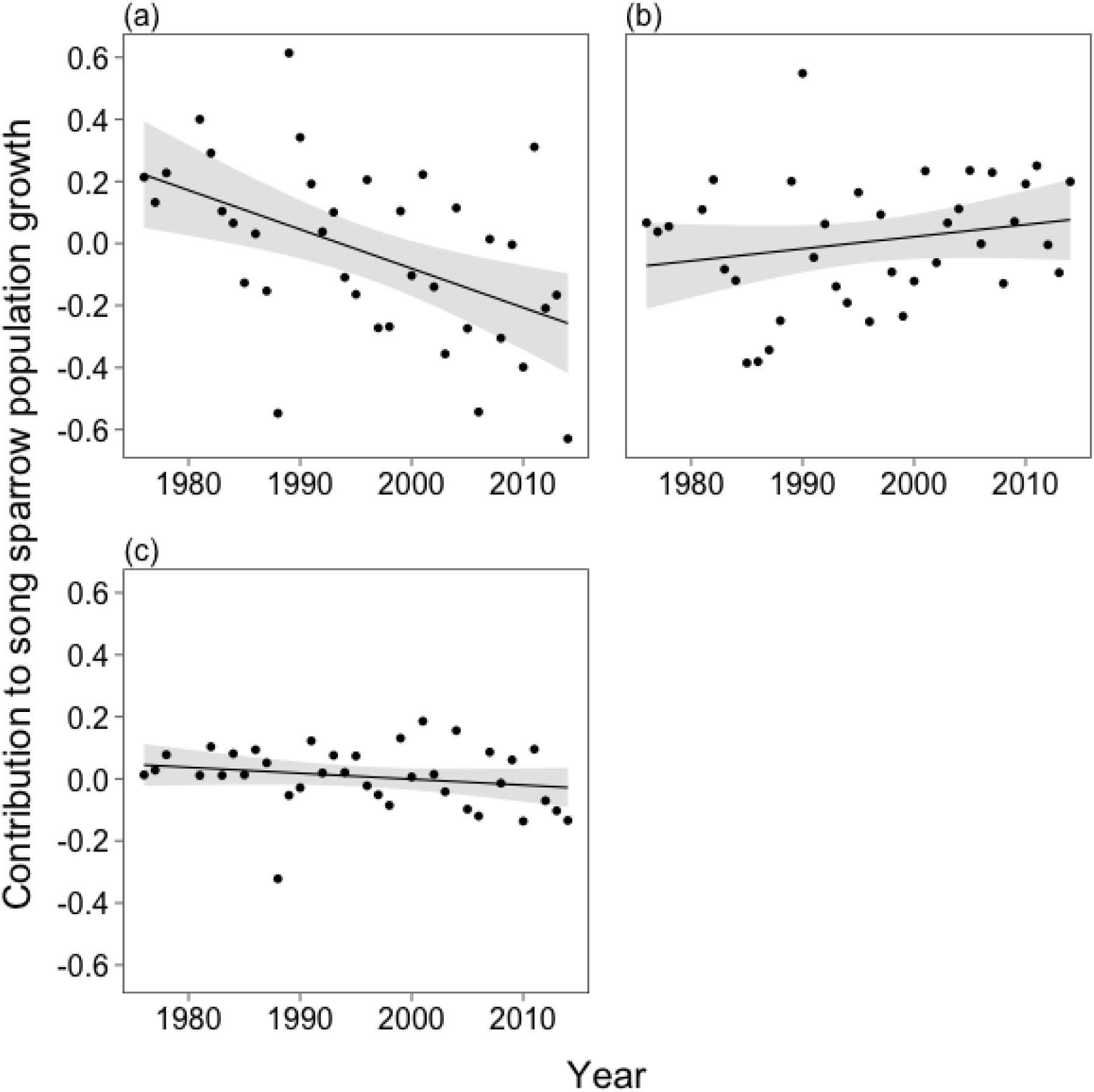
Contributions of (a) juvenile survival, (b) adult reproductive rate and (c) adult survival to song sparrow population growth from 1975–2014, derived from a stage-structured life table response experiment (see Methods). The contribution of juvenile survival changed significantly over time, but reproductive rate and adult survival remained approximately zero, indicating that the observed decline in song sparrows is best explained by the decrease in juvenile survival (Figure 3). The shaded areas around the line indicate predicted values ± 1 SE.

The fraction of juvenile song sparrows surviving overwinter also declined as fox sparrow abundance increased (figure 4), amounting to a 40% decline in the expected value of survival from 1960 to 2015 (0.39 ± 0.06 SE and 0.23 ± 0.07 SE, respectively). Juvenile song sparrow survival was also inversely related to fox sparrow population size (β = −0.009, ± 0.004 SE, *t*(8) = −2.46, p = 0.04, R^2^ = 0.44), but unrelated to song sparrow population size (*t*(8) = −0.63, p = 0.55).

**FIGURE 4.**
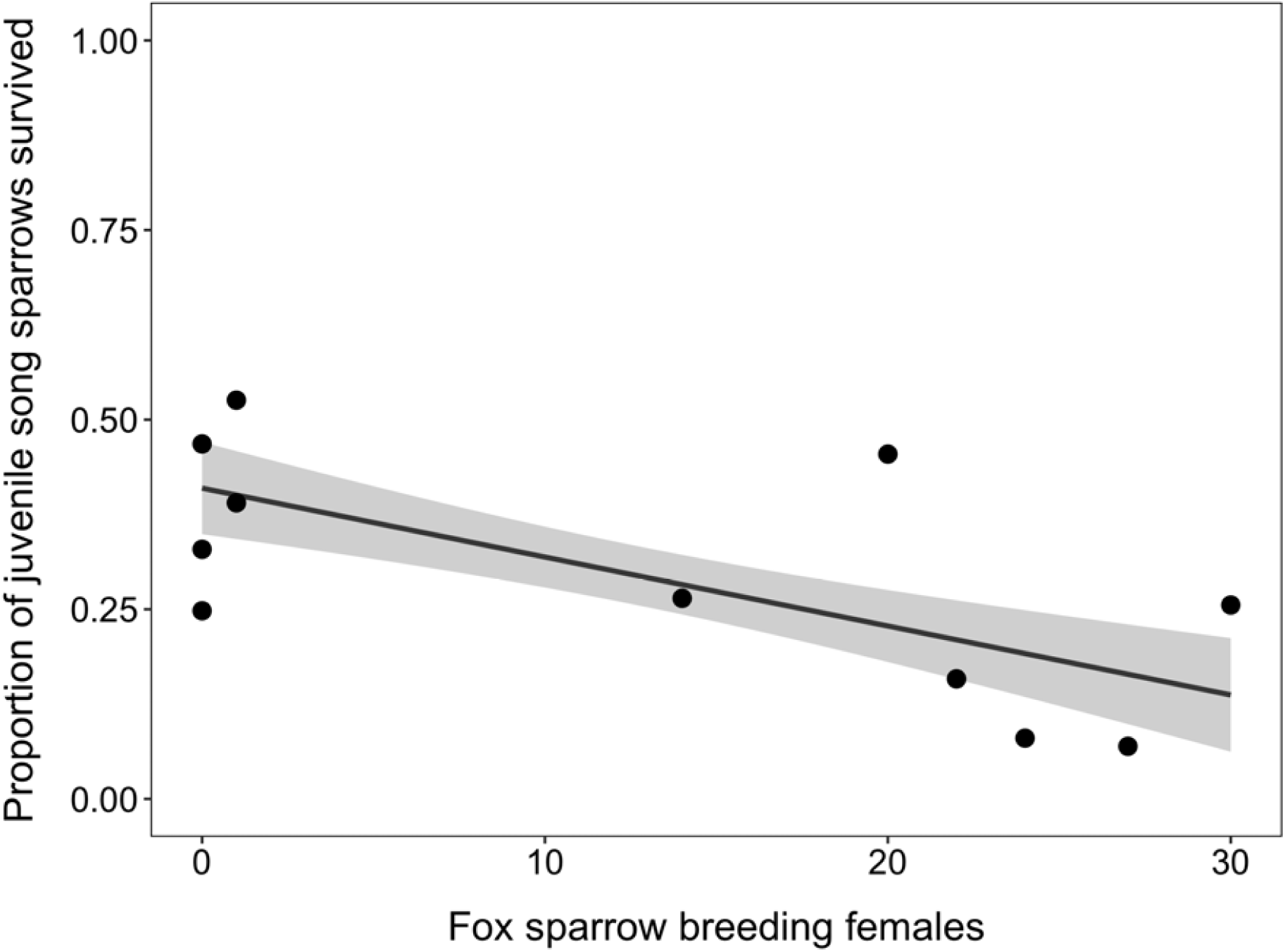
Juvenile song sparrow survival declined as the number of fox sparrow breeding females increased. The shaded areas around the line indicate predicted values ± 1 SE. The black circles are observed juvenile song sparrow survival in each study year for which fox sparrow population size was known (N_yrs_ = 11).

### Competition for winter food

Fox and song sparrows exhibited a strong preference for elderberry (mean proportion of time spent feeding 0.34 ± 0.05 SE and 0.34 ± 0.12 SE, respectively), and against blackberry (mean 0.05 ± 0.02 SE and <0.01 ± 0.003 SE, respectively). But we found no differences in the time spent feeding by each species on blackberry, Nootka rose, red elderberry or snowberry seeds (blackberry: *z*(59) = −0.81, p = 1.0, elderberry: *z*(59) = −0.19, p = 1.0, rose: *z*(59) = −0.37, p =1.0, snowberry: *z*(59) = −0.18, p = 1.0), implying complete overlap in preference for these seed species (figure 5).

**FIGURE 5.**
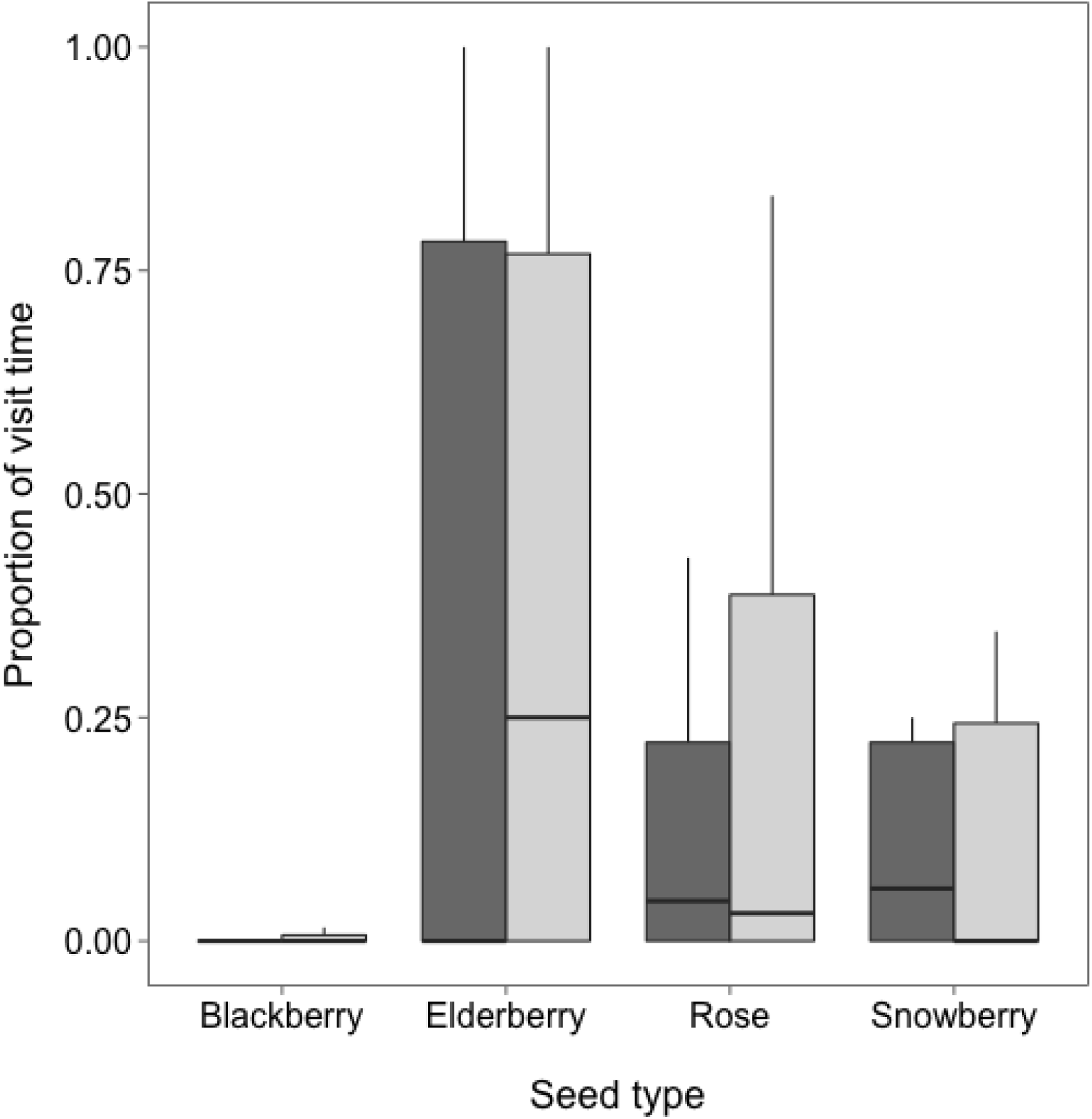
Proportion of time song (dark) and fox (light) sparrows fed on each seed type during feeder visits (see Methods). Seeds were presented by type in identical circular depressions in plywood feeders dispersed across Mandarte Island. Fox and song sparrow seed preference overlapped completely. Whiskers represent approximate 95% confidence intervals around the median (solid line), and the box spans the lower and upper quartiles (25%–75%).

Our observations of song and fox sparrows at experimental arenas baited with commercial bird seed revealed that song sparrows were displaced by fox sparrows in 91% of 68 contests (X^2^ = 25.6, df = 1, p < 0.001) in October. In a replicate experiment in early March fox sparrows displaced song sparrows in 100% of 31 interactions, obviating further analysis.

### Competition for space and nest sites

We observed no evidence of competition for space or nest sites during the breeding period. Fox and song sparrow breeding territories overlapped 100% in 2010, 2013 and 2014, and no aggressive interactions between them were observed despite regularly perching or singing within 1 m of each other. Territorial song sparrows approached conspecific mounts in simulated intrusions much more closely than fox sparrow or Swainson’s thrush mounts (*t*(41) = −7.83, p < 0.001 and *t*(41) = −8.28, p < 0.001; respectively). Song sparrows also responded similarly to fox sparrows and Swainson’s thrush mounts (*t*(41) = −0.82, p = 0.42; figure 6), indicating that song sparrows did not respond to territorial intrusions by fox sparrows as expected if these species compete for breeding space.

**FIGURE 6.**
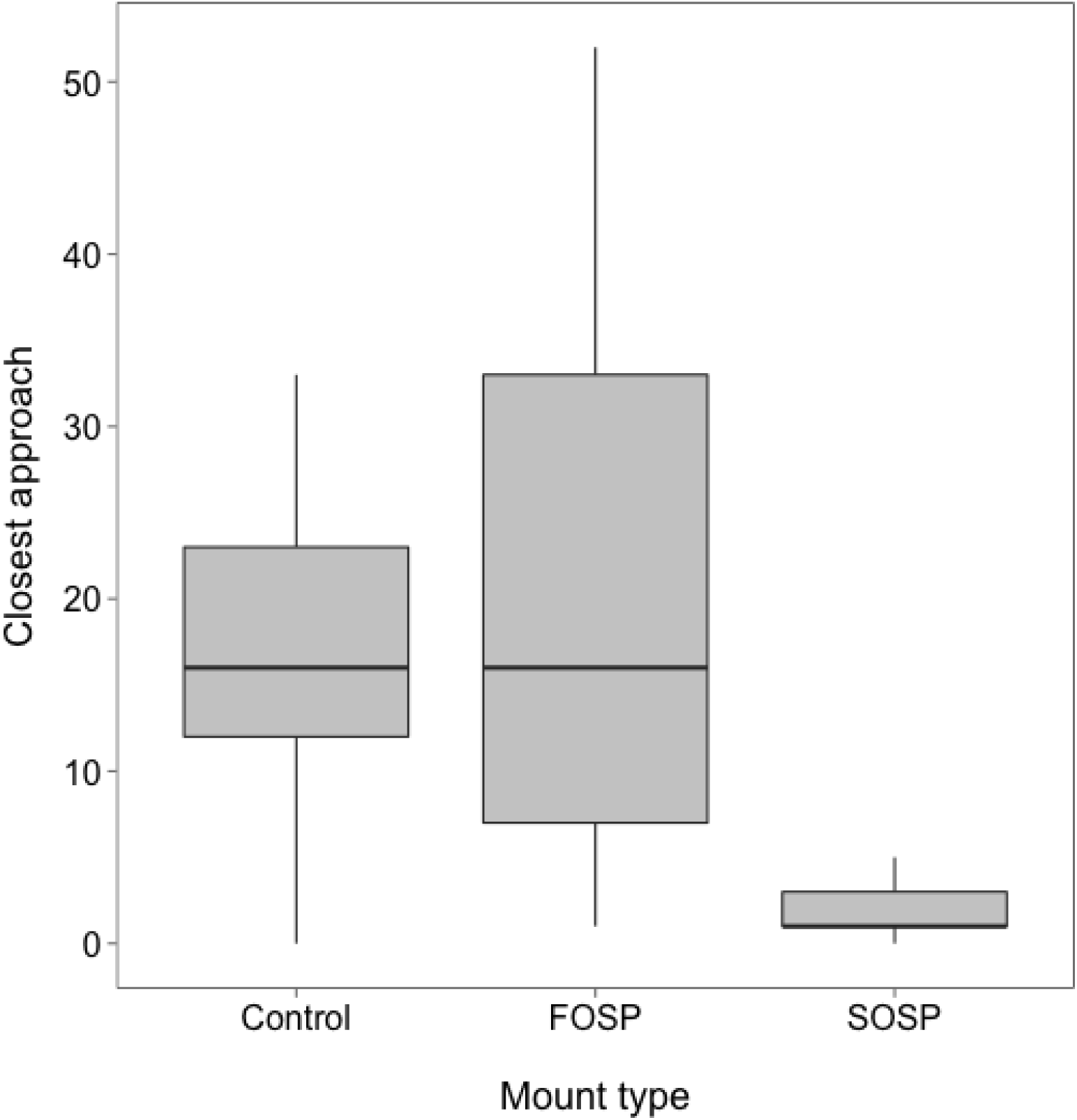
Closest approach by territorial male and female song sparrows to taxidermic mounts presented at the center of song sparrow territories during playback trials. Song sparrows (SOSP) came closer to the conspecific mount than to the fox sparrow (FOSP) or control (Swainson’s thrush) mounts, and there was no difference in song sparrow response to the fox sparrow and control mounts, indicating that song sparrows do not respond to simulated territorial intrusions by fox sparrows. Whiskers represent approximate 95% confidence intervals around the median (solid line), and the box spans the lower and upper quartiles (25%–75%).

We observed no evidence of a long-term decline in nest site quality (expected number of song sparrow young produced per nest; *t*(2671) = −1.29, p = 0.20) evaluated in 147 grid squares distributed continuously over the island and monitored annually over the study (see Methods). However, we observed a statistically significant increase in the mean annual reproductive success of female song sparrows from 1975 to 2014 (β = 0.014, SE = 0.002, *t*(643) = 6.63, p < 0.001). These results are opposite to the hypothesis that fox and song sparrows compete during the breeding period.

## DISCUSSION

We tested whether the colonization of Mandarte Is. by fox sparrows in 1975 led to the decline of the song sparrow population resident there (figure 1). Song sparrows have, on average, declined over the past 46 years whilst fox sparrows increased from 0 to 30 breeding pairs. Demographic analyses indicate that juvenile survival was the most influential of three vital rates affecting population growth in song sparrows and that it declined as fox sparrows increased (figure 4). In comparison, adult survival and reproductive rate had no detectable effect on the long-term decline in song sparrow abundance (figure 3). These findings mirror analyses conducted at much larger scales which indicate a long-term, regional decline in song sparrow abundance (Jewell and Arcese 2008), but increases in fox sparrow abundance, particularly in winter (Visty et al. 2017). Because these results are consistent with the hypothesis that interspecific competition may be contributing to song sparrow declines on Mandarte Is. and regionally, we conducted additional tests to discover potential mechanisms, focusing on competition during breeding and overwinter periods.

Contrary to the idea that fox and song sparrows compete for breeding space, we observed complete overlap in song and fox sparrow territories. Moreover, territorial song sparrows largely ignored simulated intrusions by fox sparrows (and the control), despite responding strongly to simulated intrusions by song sparrows (figure 6). These results are opposite to expectation if fox and song sparrows compete by interference for breeding space (Jankowski et al. 2010; Dhondt 2012).

We also found no evidence of exploitative competition between fox and song sparrows in the breeding period. First, annual reproductive rate in female song sparrows increased as song sparrow population size declined and fox sparrows increased (figure 3c), opposite to expectations under exploitative competition (Dhondt 2012), but consistent with earlier reports of density-dependent reproductive success in song sparrows (Arcese and Smith 1988; Arcese et al. 1992). Second, we detected no change in habitat quality for song sparrows (*cf* Germain and Arcese 2014; Crombie et al. 2017), contrary to the expectation that sharing habitat with fox sparrows might reduce food or nest site availability during breeding.

Intraspecific competition for winter food and space are well-known to affect juvenile survival and population growth in song sparrows (Nice 1943; Arcese 1989; Arcese et al. 1992; Wilson and Arcese 2008), raising the possibility that interspecific competition with fox sparrows may also reduce survival in juvenile song sparrows sufficient to cause population decline. Specifically, Dhondt (2012) notes that competitive exclusion becomes more likely when, in the presence of intraspecific competition for a limiting resource, the addition of an interspecific competitor further reduces the fitness of subordinate competitors by further reducing access to those resources. Consistent with these expectations, we observed a strong overlap in preference for native seeds in fox and song sparrows, mirroring the results of Willson (1971) who reported strong overlap in preference for commercial seed in Illinois, USA, and also found fox sparrows to be significantly more efficient at handling seeds on average. Moreover, on Mandarte Is., fox sparrows excluded song sparrows from access to supplemental food in 91 and 100% of contests in October and March, respectively. Because these periods correspond with annual peaks in intraspecific aggression and dispersal in song sparrow (Arcese 1989; Wilson and Arcese 2008), these findings suggest that fox sparrows limit song sparrow abundance on Mandate Is. via aggressive competition for winter fool. Overall, therefore, our findings are consistent with the hypothesis that range shifts in colonizing species have the potential to drive community composition via interspecific competition.

The coexistence of interspecific competitors has been shown to depend in part on the ability of species to partition resources in ways that allow each to maintain positive growth rates. In ground finches (*Geospiza* spp) on the Galapagos Islands, a drought-mediated decline in seed abundance intensified competition between a resident and colonist species, but also facilitated their rapid morphological divergence and coexistence (Grant and Grant 2006). Stuart et al. (2014) also reported the rapid evolution of feeding behavior and morphology in a native lizard following the invasion of its habitat by a competitively dominant congener. Similarly, Jankowski et al. (2010) demonstrated a strong role for interspecific competition in maintaining elevational range boundaries in congeneric Andean forest birds, but noted that as ranges shift upwards, the species at the top may be limited in their response. These studies illustrate a potential for ‘evolutionary rescue’ to facilitate co-existence via rapid adaptation (e.g., Carlson et al. 2014), and imply that song and fox sparrows are most like to co-exist where sufficient heterogeneity in habitat type or resource use allows each species to diverge sufficiently in its use of limiting resources to maintain positive growth rates.

In contrast, low habitat heterogeneity on Mandarte Is. (Lameris et al. 2016; Crombie et al. 2017), small population size and low juvenile survival rates in song sparrows all increase the likelihood of their local extirpation on Mandarte Is. (Arcese et al. 1992; Arcese and Marr 2006). Although divergence in quantitative traits potentially affecting co-existence is possible (Schluter and Smith 1986), small population size, random genetic drift and gene swamping are likely to limit rapid local adaptation in the Mandarte Is. song sparrow population (Keller et al. 2001; Marr et al. 2002). Given regional increases in fox sparrow abundance (Visty et al. 2017) and declines in song sparrow abundance (Jewell and Arcese 2008; National Audubon Society 2010; Sauer et al. 2017), it remains an open question as to how competition between these species may be affecting abundance at larger spatial scales.

Alternative explanations for the decline of our focal song sparrow population seem unlikely. Jewell and Arcese (2008) showed that brown-headed cowbirds (*Molothrus ater*) can limit song sparrow population growth by reducing reproductive success, but cowbirds were absent from Mandarte Is. in 16 of 17 yrs since 2000, and female reproductive success increased over the course of our study. Severe weather can also decimate sparrow populations (Arcese et al. 1992; Keller et al. 1994; Smith et al. 2006) but has been ameliorated by climate warming (P. Arcese and R. Norris, unpubl. res). Despite changes in vegetation cover, we detected no reduction in the cover of fruiting shrubs (Lameris et al. 2016) or increases in nest failure (Crombie et al. 2017). Overall, therefore, our results strongly support the hypothesis that fox sparrows caused the Mandarte Is. song sparrow population to decline, but more work is needed to understand how fox sparrows may limit song sparrow abundance and distribution regionally.

Shifts in species distribution could have far-reaching effects on plant and animal communities via their effects on predation, pathogens and competition (e.g., Simberloff 2005; Parmesan 2006; Early and Sax 2014; Elmhagen et al. 2015; Rodewald and Arcese 2016). Although the threat of competitive exclusion by colonizing species is sometimes downplayed (Davis 2003; Gurevitch and Padilla 2004; Krosby et al. 2015), novel competitive interactions can be expected to increase as climate and habitat change promote shifts in species ranges further. Because interspecific competition can act subtly in communities (Dhondt 2012), long-term and experimental studies of the competitive exclusion of native species by colonists undergoing range expansion will be needed to predict community dynamics in the future. Our results indicate that in the absence of ecological or evolutionary shifts in niche dimension, range expansions by dominant competitors have the potential to cause the extirpation of historically resident species when competitive interactions between them are strong and resources not equitably partitioned.

## ACKNOWLEDGEMENTS

We thank many people that have contributed to monitoring on Mandarte Is. and the Tsawout and Tseycum Bands who kindly provide us permission to work there. We are grateful to J.M. Reid, L. Keller, R. Schuster, M. Crombie, J. Krippel, N. Morrell, E. Hampshire and N. Chen for help with data, experimental design, analysis, or preparation of the manuscript. Our work was supported by the University of British Columbia, W. and H. Hesse, the American Ornithologists’ Union, and Natural Sciences and Engineering Research Council of Canada.

## LITERATURE CITED

Arcese P (1989) Intrasexual competition, mating system and natal dispersal in song sparrows. Anim Behav 38:958–979. doi: 10.1016/S0003-3472(89)80137-X

Arcese P, Marr AB (2006) Population viability in the presence and absence of cowbirds, catastrophic mortality, and immigration. In: Smith JNM, Keller LF, Marr AB, Arcese P (eds) Conservation and biology of small populations: the song sparrows of Mandarte Island. Oxford University Press, New York, New York, USA, pp 175–191

Arcese P, Smith JNM (1985) Phenotypic correlates and ecological consequences of dominance in song sparrows. J Anim Ecol 54:817–830. doi: Article

Arcese P, Smith JNM (1988) Effects of population density and supplemental food on reproduction in song sparrows. J Anim Ecol 57:119–136.

Arcese P, Smith JNM, Hochachka WM, et al (1992) Stability, regulation, and the determination of abundance in an insular song sparrow population. Ecology 73:805–822. doi: 10.2307/1940159

Arcese P, Sogge MK, Marr AB, Patten MA (2002) Song sparrow (Melospiza melodia). In: Poole A, Gill F (eds) The birds of North America, No. 704. The Birds of North America Inc., Philadelphia, PA, USA,

Bennett JR, Dunwiddie PW, Giblin DE, Arcese P (2012) Native versus exotic community patterns across three scales: Roles of competition, environment and incomplete invasion. Perspect Plant Ecol Evol Syst 14:381–392. doi: 10.1016/j.ppees.2012.10.001

Carlson SM, Cunningham CJ, Westley PAH (2014) Evolutionary rescue in a changing world. Trends Ecol Evol 29:521–530. doi: 10.1016/j.tree.2014.06.005

Caswell H (1996) Analysis of life table response experiments II. alternative parameterizations for size- and stage-structured models. Ecol Modell 88:73–82. doi: 10.1016/0304-3800(95)00070-4

Chesson P (2000) Mechanisms of maintenance of species diversity. Annu Rev Ecol Syst 31:343–366. doi: 10.1146/annurev.ecolsys.31.1.343

Crombie MD, Germain RR, Arcese P (2017) Nest-site preference and reproductive performance of Song Sparrows (Melospiza melodia) in historically extant and colonist shrub species. Can J Zool 95:115–121. doi: 10.1139/cjz-2016-0189

Davies KE, Chesson P, Harrison S, et al (2005) Spatial heterogeneity explains the scale dependence of the native-exotic diversity relationship. Ecology 86:1602–1610.

Davis MA (2003) Biotic globalization: does competition from introduced species threaten biodiversity? Bioscience 53:481. doi: 10.1641/0006-3568(2003)053[0481:BGDCFI]2.0.CO;2

Davis MB, Shaw RG (2001) Range shifts and adaptive responses to quaternary climate change. Science 292:673–679.

de Kroon H, Plaisier A, Groenendael J Van, Caswell H (1986) Elasticity: the relative contribution of demographic parameters to population growth rate. Ecology 67:1427–1431.

Dhondt AA (2012) Interspecific competition in birds. Oxford University Press, New York, New York, USA

Doherty TS, Glen AS, Nimmo DG, et al (2016) Invasive predators and global biodiversity loss. Proc Natl Acad Sci U S A 113:11261–11265. doi: 10.1073/pnas.1602480113

Drent PJ, van Tets GF, Tompa FS, Vermeer K (1964) The breeding birds of Mandarte Island, British Columbia. Can Field-Naturalist 78:208–263.

Early R, Sax DF (2014) Climatic niche shifts between species' native and naturalized ranges raise concern for ecological forecasts during invasions and climate change. Glob Ecol Biogeogr 23:1356–1365. doi: 10.1111/geb.12208

Elmhagen B, Eriksson O, Lindborg R (2015) Implications of climate and land-use change for landscape processes, biodiversity, ecosystem services, and governance. Ambio 44 Suppl 1:S1–5. doi: 10.1007/s13280-014-0596-6 ESRI (2011) ArcGIS Desktop: Release 10.

Germain RR, Arcese P (2014) Distinguishing individual quality from habitat preference and quality in a territorial passerine. Ecology 95:436–445. doi: 10.1890/13-0467.1

Germain RR, Schuster R, Delmore KE, Arcese P (2015) Habitat preference facilitates successful early breeding in an open-cup nesting songbird. Funct Ecol 29:1522–1532. doi: 10.1111/1365-2435.12461

Grant PR, Grant BR (2006) Evolution of character displacement in Darwin's finches. Science 313:224–226. doi: 10.1002/9780470114735.hawley00624

Gurevitch J, Padilla DK (2004) Are invasive species a major cause of extinctions? Trends Ecol Evol 19:470–474. doi: 10.1016/j.tree.2004.07.005

Hothorn T, Bretz F, Westfall P (2008) Simultaneous inference in general parametric models. Biometrical J 50:346–363.

Jankowski JE, Robinson SK, Levey DJ (2010) Squeezed at the top: interspecific aggression may constrain elevational ranges in tropical birds. Ecology 91:1877–1884.

Jewell KJ, Arcese P (2008) Consequences of parasite invasion and land use on the spatial dynamics of host populations. J Appl Ecol 45:1180–1188. doi: 10.1111/j.1365-2664.2008.01503.x

Keller LF, Arcese P, Smith JNM, et al (1994) Selection against inbred song sparrows during a natural population bottleneck. Nature 372:356–357.

Keller LF, Jeffery KJ, Arcese P, et al (2001) Immigration and the ephemerality of a natural population bottleneck: evidence from molecular markers. Proc Royal Soc Biol 268:1387–1394.

Kramer CY (1956) Extension of multiple range tests to group means with unequal number of replications. Biometrics 12:307–310.

Krosby M, Wilsey CB, Mcguire JL, et al (2015) Climate-induced range overlap among closely related species. Nat Clim Chang 5:883–886. doi: 10.1038/NCLIMATE2699

Lameris TK, Bennett JR, Blight LK, et al (2016) A century of ecosystem change: human and seabird impacts on plant species extirpation and invasion on islands. PeerJ 4:e2208. doi: 10.7717/peerj.2208

Loarie SR, Carter BE, Hayhoe K, et al (2008) Climate change and the future of California's endemic flora. PLoS One 3:e2502. doi: 10.1371/journal.pone.0002502

MacDougall AS, Gilbert B, Levine JM (2009) Plant invasions and the niche. J Ecol 97:609–615. doi: 10.1111/j.1365-2745.2009.01514.x

Marr AB, Keller LF, Arcese P (2002) Heterosis and outbreeding depression in descendants of natural immigrants to an inbred population of song sparrows (melospiza melodia). Evolution 56:131–142. doi: 10.1111/j.0014-3820.2002.tb00855.x

Melbourne BA, Cornell H V., Davies KF, et al (2007) Invasion in a heterogeneous world: resistance, coexistence or hostile takeover? Ecol Lett 10:77–94. doi: 10.1111/j.1461-0248.2006.00987.x

National Audubon Society (2010) The Christmas bird count historical results [Online]. http://www.christmasbirdcount.org. Accessed 16 Jun 2015

Nice MM (1943) Studies in the life history of the song sparrow. II. The behavior of the song sparrow and other passerines. New York

Parmesan C (2006) Ecological and evolutionary responses to recent climate change. Annu Rev Ecol Syst 37:637–669.

R Core Team (2015) R: A language and environment for statistical computing.

Reaser JK, Meyerson LA, Cronk Q, et al (2007) Ecological and socioeconomic impacts of invasive alien species in island ecosystems. Environ Conserv 34:98. doi: 10.1017/S0376892907003815

Rodewald AD, Arcese P (2016) Direct and Indirect Interactions between Landscape Structure and Invasive or Overabundant Species. Curr Landsc Ecol Reports 1:30–39. doi: 10.1007/s40823-016-0004-y

Sauer JR, Niven DK, Hines JE, et al (2017) The North American Breeding Bird Survey, Results and Analysis 1966 - 2015. Version 2.07.2017 USGS Patuxent Wildlife Research Center,Laurel, MD.

Sax DF, Stachowicz JJ, Brown JH, et al (2007) Ecological and evolutionary insights from species invasions. Trends Ecol Evol 22:465–471. doi: 10.1016/j.tree.2007.06.009

Schluter D, Smith JNM (1986) Natural Selection on Beak and Body Size in the Song Sparrow. Evolution 40:221. doi: 10.2307/2408803

Shea K, Chesson P (2002) Community ecology theory as a framework for biological invasions. Trends Ecol Evol 17:170–176.

Simberloff D (2005) Non-native species DO threaten the natural environment! J Agric Environ Ethics 18:595–607. doi: 10.1007/s10806-005-2851-0

Smith JNM, Marr AB, Arcese P, Hochachka WM (2006) Fluctuations in numbers: population regulation and catastrophic mortality. In: Smith JNM, Keller LF, Marr AB, Arcese P (eds) Conservation and biology of small populations: the song sparrows of Mandarte Island. Oxford University Press, New York, New York, USA, pp 43–64

Sorte CJB, Williams SL, Carlton JT (2010) Marine range shifts and species introductions: comparative spread rates and community impacts. Glob Ecol Biogeogr 19:303–316. doi: 10.1111/j.1466-8238.2009.00519.x

Stuart YE, Campbell TS, Hohenlohe PA, et al (2014) Rapid evolution of a native species following invasion by a congener. Science 346:463–466. doi: 10.1126/science.1257008

Stubben C, Milligan B (2007) Estimating and analyzing demographic models using the popbio package in R. J Stat Softw 22:1–23.

Tompa FS (1963) Factors determining the numbers of song sparrows on Mandarte Island, B.C. University of British Columbia

Tukey J (1949) Comparing individual means in the analysis of variance. Biometrics 5:99–114.

Visty H, Wilson S, Germain RR, et al (2017) Demography of sooty fox sparrows (Passerella iliaca unalaschcensis) following a shift from a migratory to resident life history. Can J Zool. In Press.

Weckstein JD, Kroodsma DE, Faucett RC (2002) Fox sparrow (Passerella iliaca). In: Poole A (ed) The birds of North America online. Cornell Lab of Ornithology, Ithaca, New York, USA.

Willson MF (1971) Seed selection in some North American finches. Condor 73:415–429.

Wilson AG, Arcese P (2008) Influential factors for natal dispersal in an avian island metapopulation. J Avian Biol 39:341–347. doi: 10.1111/j.0908-8857.2008.04239.x

Wilson S, Norris DR, Wilson AG, Arcese P (2007) Breeding experience and population density affect the ability of a songbird to respond to future climate variation. Proc R Soc B-Biological Sci 274:2539–2545. doi: 10.1098/rspb.2007.0643

